# *Ureaplasma parvum* SMC-ScpAB complex is capable of loop extrusion and demonstrates properties that distinguish it from *Bacillus subtilis* homologue

**DOI:** 10.1101/2025.05.11.653314

**Authors:** Natalia A. Rumyantseva, Antonina P. Sapozhnikova, Aizilya A. Khasanova, Ekaterina Yu. Zapryagaeva, Maria A. Kudryavtseva, Natalia E. Morozova, Alexey D. Vedyaykin

## Abstract

Structural Maintenance of Chromosomes (SMC) complexes are present in virtually all organisms and perform a variety of functions associated with maintaining the integrity and spatial organization of DNA. The best-studied SMC complexes are eukaryotic condensins, cohesins, and Smc5/Smc6. It is extremely important that eukaryotic SMC have been shown to exhibit the ability for so-called loop extrusion *in vitro*, which is the active formation of loops from DNA molecules and is a cosequence of the DNA translocase activity of SMC complexes. For majority of bacterial SMC complexes, including the most widespread Smc-ScpAB complex, loop extrusion has not yet been demonstrated *in vitro*, although it is in good agreement with the results of *in vivo* experiments. In this work, we compared the properties of two Smc-ScpAB complexes from different organisms, *Bacillus subtilis* and *Ureaplasma parvum*. The results of the work indicate significant differences in the properties of these homologous complexes. In particular, the Smc-ScpAB complex of *U. parvum* was shown to have the ability to extrude loops, which was not observed for *B. subtilis* SMC.

## Introduction

Structural maintenance of chromosomes (SMC) complexes are large protein structures that interact with DNA and participate in key processes, including DNA replication and segregation, DNA repair, DNA compaction, protection aginst foreign DNA, etc. [1]. SMC complexes are probably present in all domains of living organisms, although in limited cases they are not essential. For example, in a number of bacteria the genes encoding proteins of the SMC complex can be deleted, although this leads to defects in DNA segregation, especially under conditions of rapid growth [2]. However, even in mollicutes, some of which are close to the so-called “minimal” cell, SMC complexes have been found, which confirms the extreme importance of these complexes and probably indicates their essentiality for at least some living organisms [3]. SMC complexes typically consist of three main components: ATPase, kleisin, and accessory protein(s) (HAWK or KITE) [4]. In all complexes, the ATPase is a dimer (a heterodimer in eukaryotes or a homodimer in bacteria) consisting of two monomer arms. Each monomer consists of two globular domains (hinge and head domains) connected by a long (about 50 nm) coiled coil linker. One of the globular domains (a hinge) ensures the dimerization of the monomers, which leads to the formation of a characteristic V-shaped structure. Other globular domain, the head domain, is also capable of interacting with the neighboring head domain, and the interaction depends on ATP binding. In addition to the ATPase dimer, the complex contains a kleisin, which interacts with the head domains of the ATPase and promotes the formation of a closed ring structure of the complex, as well as an auxiliary protein HAWK (only in eukaryotes) or KITE (found in both eukaryotes and prokaryotes), which plays a regulatory role. In bacteria, one of the most common SMC complexes is Smc-ScpAB, in which Smc is an ATPase, ScpA is a kleisin, and ScpB is a KITE protein [5]. Currently, the consensus model of SMC action is the loop extrusion, in which SMC complexes with ATP-dependent DNA translocase activity, are capable of forming loops from DNA molecules. This model is supported by a large amount of data obtained in a wide range of organisms using various methods, including the analysis of the spatial organization of genomes using the Hi-C method [6, 7]. Loop extrusion has now been directly demonstrated for eukaryotic SMC complexes – condensins [8], cohesins [9] and Smc5/Smc6 complexes [10] — *in vitro* at the single-molecule level. At the same time, for bacterial SMC complexes, there are currently only limited data confirming loop extrusion obtained at the single-molecule level – only for the *E. coli* MukBEF complex [11] И *P. aeruginosa* Wadjet [12]. At the same time, for the most widespread Smc-ScpAB complex, such data are absent to the date. In this work, the properties of Smc-ScpAB complexes from two evolutionarily distant bacteria, *Ureaplasma parvum* and *Bacillus subtilis*, were investigated. Several important differences between these complexes in their ability to interact with DNA and ATPase activity were revealed. The ability of the Smc-ScpAB complex of *U. parvum* to extrude loops was demonstrated for the first time.

## Materials and Methods

### 1. Purification of proteins

The genes encoding the proteins of the Smc-ScpAB complexes of *U. parvum* and *B. subtilis* were cloned into the pET21a vector using standard methods in such a way that the His-tag of the produced proteins was located at the C-terminus, similar to the Smc gene of *U. parvum* which was previously cloned [13].

Proteins were produced in *E. coli* cells of the Rosetta (DE3) strain. The cells were pre-cultured for 2 h, then the expression was induced by adding IPTG to 1 mM. Proteins were produced at 18°C overnight. The culture was centrifuged and stored at -80C.

Cell pellets were resuspended in a buffer of appropriate composition prior to purification, with the exception of *U. parvum* ScpA, the cell pellet of which was resuspended prior to freezing in buffer (20 mM Tris-HCl pH = 8, 20% glycerol, 1 mM EDTA, 1 mM DTT, 0.05% Tween-20) and frozen in the buffer. For *B. subtilis* Smc, the following buffer was used for resuspension: 20 mM Tris-HCl, 400 mM Ammonium Sulfate, 25 mM Imidasole, pH 7.5. The cells were lysed by sonication with the addition of lysozyme. After sonication, the lysate was clarified by centrifugation and filtration. Then, for preliminary purification from DNA, the clarified lysate was passed through a HiTrap Q anion exchange column, the flow-through was collected, which was then applied to a column with Ni-NTA resin. Elution was carried out by stepwise increasing the concentration of imidazole. Eluates were frozen in liquid nitrogen and stored at -80°C with the addition of glycerol to 20%, NaCl to 100 mM and DTT to 0.1 mM.

For *B. subtilis* ScpA and ScpB, the following buffer was used for resuspension: 20 mM Tris, pH=7.5, 500 mM NaCl. They were purified using Ni-NTA resin similarly to Smc, then diluted tenfold in a salt-free buffer (50 mM Tris, pH=7.5, 1 mM DTT) and applied to a MonoQ column with anion exchange resin, after which they were eluted by increasing the concentration of NaCl. They were dialyzed into storage buffer (50 mM Tris, pH=7.5, 1 mM EDTA, 0.1 mM DTT, 100 mM NaCl + 20% Glycerol), frozen in liquid nitrogen and stored at -80°C.

*U. parvum* Smc protein was isolated similarly to the method described in the article [13].

The *U. parvum* ScpA protein was isolated similarly to Smc.

The *U. parvum* ScpB protein was isolated similarly to ScpB of *B. subtilis*. All proteins were concentrated by dialysis against storage buffer with the addition of 1 M sucrose, since the use of diafiltration resulted in precipitation of proteins.

### 2. Analysis of protein-protein interactions in SMC complexes

Protein-protein interactions were analyzed by cross-linking protein complexes with formaldehyde and subsequent analysis by SDS-PAGE. For this purpose, proteins were first dialyzed against the buffer: Hepes 50 mM (pH = 7.5), NaCl 100 mM, DTT 0.1 mM, 20% Glycerol. Then, proteins in various combinations were incubated in the buffer indicated above with or without the addition of formaldehyde (0.4%). Then, samples were denatured at 65°C with the addition of SDS and BME and analyzed using the standard SDS-PAGE method. Mass spectrometry (peptide fingerprinting, similar to that indicated in the article[13]) was used to identify proteins.

### 3. SMC-DNA interaction assay (EMSA)

Circular plasmid pUC19 dual (9396 bp) was used as DNA substrate [14]. The reaction mixture was based on buffer (10 mM Tris-HCl, pH=7.0, 50 mM NaCl, 50 mM KCl, 1 mM DTT, 5 mM MgCl_2_, 2.5 mM ATP), supplemented (where necessary) with 10 pM of each of the proteins of SMC complexes (except *B. subtilis* Smc – 5 pM), and also 50 ng of DNA. Samples were loaded on 0.7% agarose gel containing ethidium bromide (1/10000) for DNA visualization, in Tris-acetate buffer, the electrophoresis was performed, and then visualized using ChemiDoc.

### 4. Determination of ATPase activity of SMC complexes

The reaction mixture consisted of 10 mM Tris-HCl, pH 7.0, 50 mM NaCl, 50 mM KCl, 1 mM DTT, 5 mM MgCl_2_, 2.5 mM ATP, and (where necessary) each of the SMC complex proteins (to a final concentration of 100 nM), and 50 ng DNA. The mixture was incubated at 37°C for specified time intervals up to 1 hour. Then 10 μl of the reaction mixture was transferred to a 96-well flat-bottomed plate. 100 μl of color reagent (consisting of 1 part 4.2% ammonium molybdate, 3 parts 0.045% malachite green solution, and 0.1% Triton-X100) were added to the samples and mixed. The absorbance of the solutions at 630 nm was analyzed using a CLARIOstar spectrophotometer. A calibration curve corresponding to standards with known free phosphate concentrations was used to determine the concentration of hydrolyzed ATP in the samples.

### 5. Visualization of loop extrusion

Coverslips for microscopy were cleaned using plasma in air atmosphere. After that, they were treated with APTES ((3-Aminopropyl) triethoxysilane) solution and Biotin-Polyethylene Glycol (mPEG + Biotin-PEG + bicarbonate buffer) solution. Modification of λ phage DNA ends was carried out by adding biotinylated dCTP using Klenow fragment. To immobilize DNA on biotinylated coverslips, a flow chamber was formed consisting of a slide and a coverslip, where the modified surface of the coverslip was oriented toward the slide. The chamber was washed with PBS, then streptavidin solution was added to it. After that, bacteriophage λ DNA modified with biotin at both ends was added to the chamber. After that, the flow chamber was filled with a buffer containing Tris-HCl (pH=7) - 10 mM, NaCl - 50 mM, KCl - 50 mM, MgCl_2_ - 5 mM, MEA - 100 mM, PCA - 2.5 uM, PCD - 0.1 U/ml, COT - 2 mM, ATP - 1 mM, YOYO-1 - 200 nM. To straighten the DNA, a solution flow was created in the chamber by microfluidics. As a result, the DNA was fixed to the glass by its ends, in some cases forming a characteristic U-shaped structure in the flow. This process was visualized using fluorescence microscopy in the YOYO-1 dye channel.

## Results and Discussion

### 1. Purified Smc, ScpA and ScpB Proteins from *B. subtilis and U. parvum* are Capable to Form SMC complexes that demonstrate distinct interactions

Individual proteins (Smc, ScpA and ScpB) of the SMC complexes of *B. subtilis* and *U. parvum* were purified using metal affinity and anion exchange chromatography. All 6 proteins, when analyzed by SDS-PAGE, demonstrate a major band corresponding in size to the expected mass of the protein (*B. subtilis*: Smc – 135.5 kDa, ScpA – 31 kDa, ScpB – 23.2 kDa; *U. parvum*: Smc – 111.7 kDa, ScpA – 34.6 kDa, ScpB – 41.8 kDa), as well as a certain number of minor bands corresponding to impurity proteins (see Fig. 1a and 1c, the first 3 bands to the left starting from the marker correspond to purified proteins). However, the ScpB protein of *U. parvum* demonstrates lower purity than the other obtained proteins, since the impurity bands are much more noticeable than in the case of other proteins. All impurity bands were analyzed by mass spectrometry to identify impurities, among them no proteins binding to DNA or possessing ATPase activity were found. Cross-linking of the complexes using formaldehyde allowed observing oligomerization and interaction of proteins in SMC complexes with each other (see Fig. 1). The ability of Smc and ScpB to dimerize and direct interactions between proteins ScpA and ScpB as well as Smc and ScpA of *B. subtilis* were confirmed. These data are fully consistent with published data [15, 16], which indicates the adequacy of the technique used. In the case of the *U. parvum* ScpA and ScpB, it was not possible to observe their interactions with each other, unlike *B. subtilis* ScpA and ScpB. Furthermore, *U. parvum* ScpB does not demonstrate visible dimerization. It is interesting to note that, unlike *B. subtilis* Smc, *U. parvum* Smc protein forms not only a dimer, but also (presumably) a tetramer. Thus, the interactions within the *U. parvum* SMC complex differ significantly from those for the *B. subtilis* SMC complex. These observations may indicate a different compositions of the complete SMC complexes (in particular, probably a different stoichiometry of the components of the complex), which requires additional study in the future. For the *B. subtilis* SMC complex, the stoichiometry of the Smc:ScpA:ScpB components is estimated differently [17], for example, 2:1:2 or 2:2:4 or 4:2:4, where Smc tetramerization (dimer of dimers) is considered to be associated with interactions of ScpA and/or ScpB proteins [2]. These differences between the studied SMC complexes may explain the differences between these complexes in ATPase activity and the ability to interact with DNA (see below).

**Figure 1.**
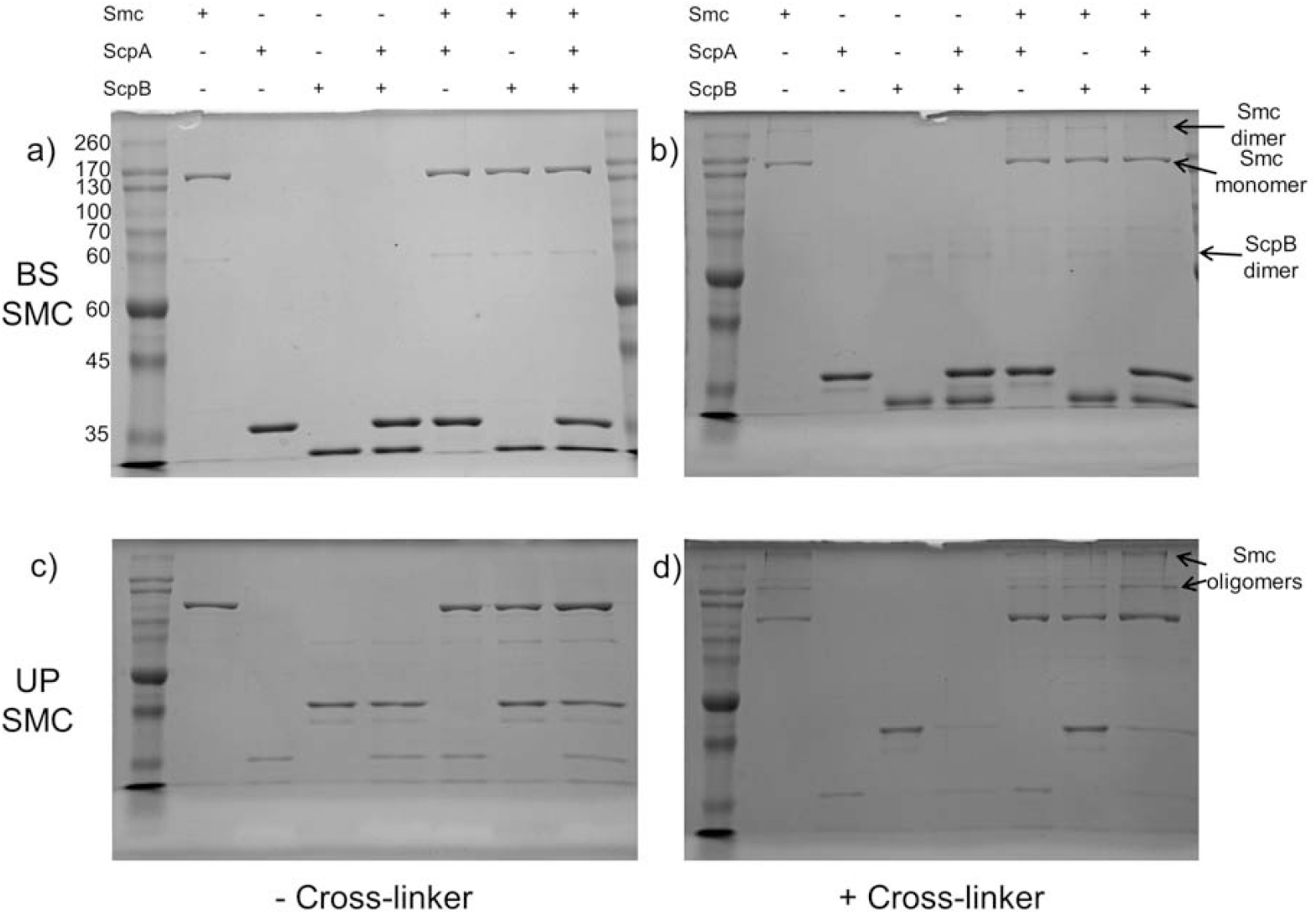
Individual proteins and their combinations of the SMC complexes of *B. subtilis* (a, b) and *U. parvum* (c, d) visualized by SDS-PAGE. The proteins without the cross-linker are shown on the left, and with the addition of the cross-linker (0.4% formaldehyde) on the right. The table on the top indicates which proteins were added to a given sample and refers to both upper and lower parts of the figure. The arrows indicate the monomers and oligomers of the Smc and ScpB proteins of *B. subtilis* and *U. parvum*

### 2. *B. subtilis and U. parvum* SMC complexes demonstrate different DNA-binding properties

We have previously established that *U. parvum* Smc protein exhibits ATP-stimulated ability to form complexes with DNA [13]. Under similar conditions, this protein behaves similarly to the *B. subtilis* Smc protein, except that the latter exhibits a significantly lower ability to bind to a linear double-stranded DNA molecule than *U. parvum* Smc (data not shown). In this work, we investigated the effect of the ScpA and ScpB proteins on the interaction of the *B. subtilis* and *U. parvum* SMC complexes with DNA, which made it possible to identify both the similarity and some differences between these complexes (see Fig. 2). Both complexes are characterized by a weakening of the interaction with the DNA molecule due to the addition of ScpA and ScpB together or separately to a sample containing Smc (as follows from an increase in the proportion of free DNA compared to a sample containing only Smc). However, unlike their *B. subtilis* homologues, *U. parvum* ScpA and ScpB demonstrate intrinsic binding to DNA (see Fig. 2b). The latter property is not characteristic of Smc-ScpAB complexes: according to published data, ScpAB proteins affect the binding of Smc to DNA, but do not independently bind to DNA. At the same time, DNA binding has been shown for the Cnd2-Cnd3 subcomplex (yeast homologue of the ScpAB subcomplex from condensin) [18, 19], although both subunits are required for DNA binding in this case.

**Figure 2.**
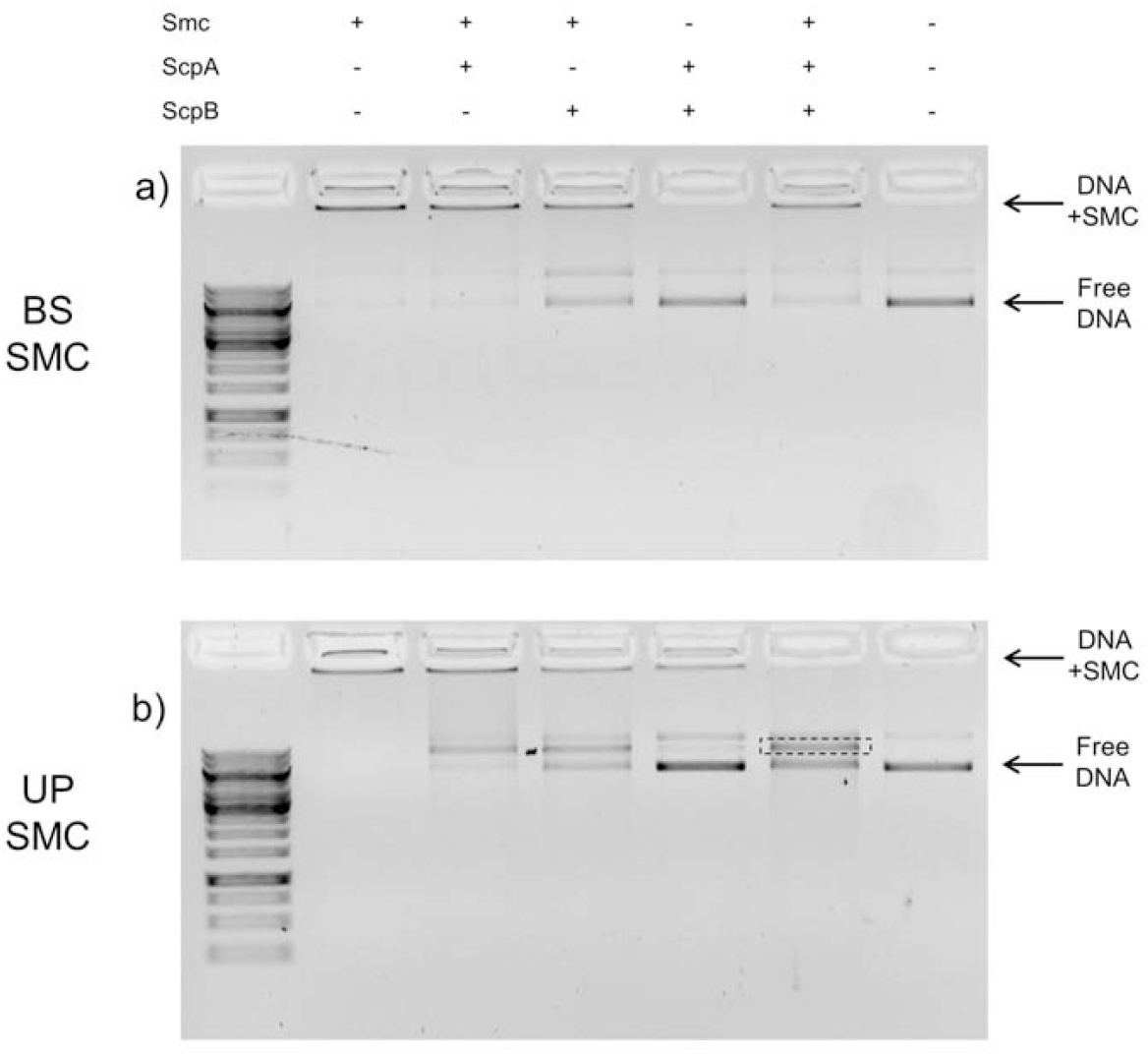
Interaction of *B. subtilis* (a) and *U. parvum* (b) SMC complexes with circular double-stranded DNA visualized by EMSA. Arrows indicate free DNA (Free DNA) and DNA in complex with proteins of the SMC complex immobilized in the well of the agarose gel (DNA+SMC). The rectangle indicates the *U. parvum* SMC-ScpAB complex with DNA. The leftmost lane shows the DNA length marker (GeneRuler 1 kb DNA Ladder)

Moreover, upon formation of the entire *U. parvum* Smc-ScpAB complex, the nature of the DNA mobility shift changes: if in the case of the *B. subtilis* Smc-ScpAB complex only 2 main DNA bands are visible – immobilized in the well (SMC and DNA complex) and free DNA, then in the case of the *U. parvum* ScpA Smc-ScpAB complex an additional third band appears (outlined in a rectangle in Fig. 2b), also corresponding to the SMC complex with DNA. A possible explanation for this observation is that the *U. parvum* partial SMC complex (as well as the partial and complete *B. subtilis* SMC complex) provides the cross-linking of several DNA molecules (which leads to the immobilization of SMC+DNA complexes in the gel well), whereas the complete *U. parvum* Smc-ScpAB complex demonstrates binding to one DNA molecule, affecting its mobility, which leads to the formation of a relatively sharm band on the gel.

### 3. *B. subtilis and U. parvum* SMC complexes demonstrate different effect of ScpAB on ATP hydrolysis

We have previously found that *U. parvum* Smc exhibits DNA-stimulated ATPase activity, which was estimated at 3.8 ATP molecules per minute per 1 Smc monomer (or 7.8 ATP molecules per minute per 1 Smc dimer) [13]. This activity is relatively low compared to *B. subtilis* Smc [20] and is comparable to that of MukB [21]. In the current work, we have shown that ScpAB significantly (several-fold) increases the ATPase activity of *U. parvum* Smc, in contrast to *B. subtilis* Smc, which exhibits high ATPase activity even in the absence of ScpAB and DNA (see Fig. 3). The ScpA and ScpB proteins of both studied complexes do not exhibit significant ATPase activity by themselves. It is interesting to note that the addition of ScpA increases the ATPase activity of *U. parvum* Smc in approximately the same way as ScpAB, whereas ScpB alone does not activate this activity. The published data on the effect of ScpAB on the ATPase activity of Smc are contradictory. For example, in the paper [20] ScpAB was shown to reduce the ATPase activity of *B. subtilis* Smc, which contradicts the results of our current work. At the same time, in another paper [17] it has been demonstrated that ScpAB increases the ATPase activity of *Geobacillus stearothermophilus* Smc by approximately 2 times. It is interesting to note that for MukBEF, a homologue of Smc-ScpAB in a number of gram-negative bacteria, the stimulation of ATPase activity by the MukEF subcomplex (a homologue of ScpAB) has also been shown [22]. Apparently, even homologous SMC complexes in different bacteria exhibit significantly different properties, including the effect of ScpAB on the ATPase activity of Smc. Thus, the ATPase activity of the *U. parvum* SMC complex is activated by both DNA and the ScpAB subcomplex, with ScpA activating the ATPase activity in the absence of ScpB. Probably, such behavior of *U. parvum* SMC complex is explained by the fact that binding of Smc, which is in a closed conformation (in which the head domains contact each other and bind but do not hydrolyze ATP), to DNA and/or ScpA stimulates its transition to an open conformation and, accordingly, ATP hydrolysis. This assumption is consistent with the above observation that ScpAB inhibits the formation of Smc complexes with DNA.

**Figure 3.**
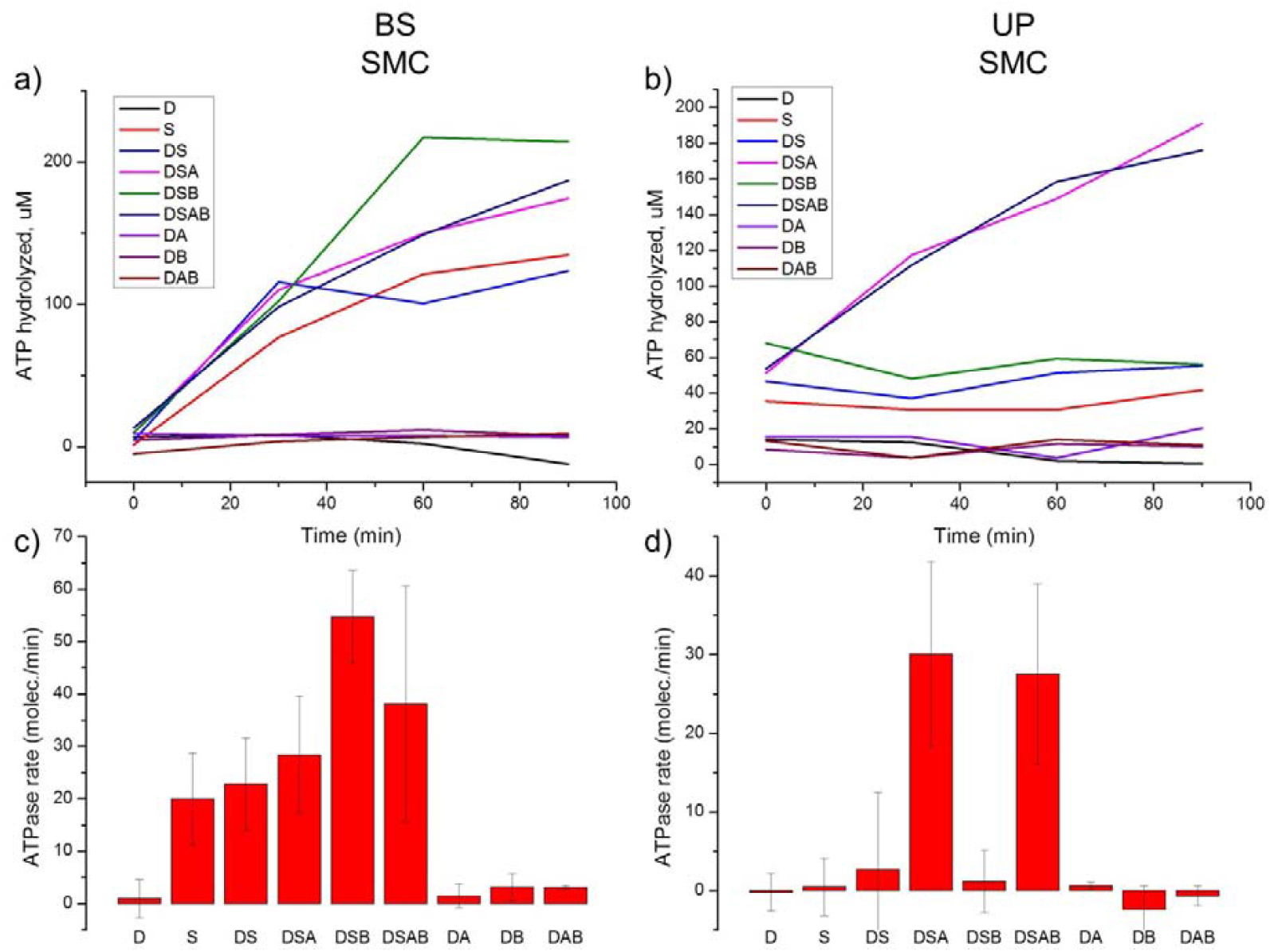
ATPase activity of proteins of the SMC complexes of *B. subtilis* (a, c) and *U. parvum* (b, d). At the top (a and b) are examples of time dependences of the concentration of hydrolyzed ATP in the presence or absence of various proteins of the SMC complexes (S – Smc, A – ScpA, B – ScpB) and DNA (D). D – in the presence of DNA and in the absence of proteins; S – in the presence of Smc and in the absence of DNA; DS – in the presence of DNA and Smc; DSA – in the presence of DNA, Smc and ScpA; DSB – in the presence of DNA, Smc and ScpB; DSAB – in the presence of DNA, Smc, ScpA and ScpB; DA – in the presence of DNA and ScpA; DB – in the presence of DNA and ScpB; DAB – in the presence of DNA, ScpA and ScpB. At the bottom (c and d) are bar graphs showing the median values of ATPase activity levels for the same combinations of proteins and DNA as in a, b. Whiskers on the graphs represent standard deviations

**Figure 4.**
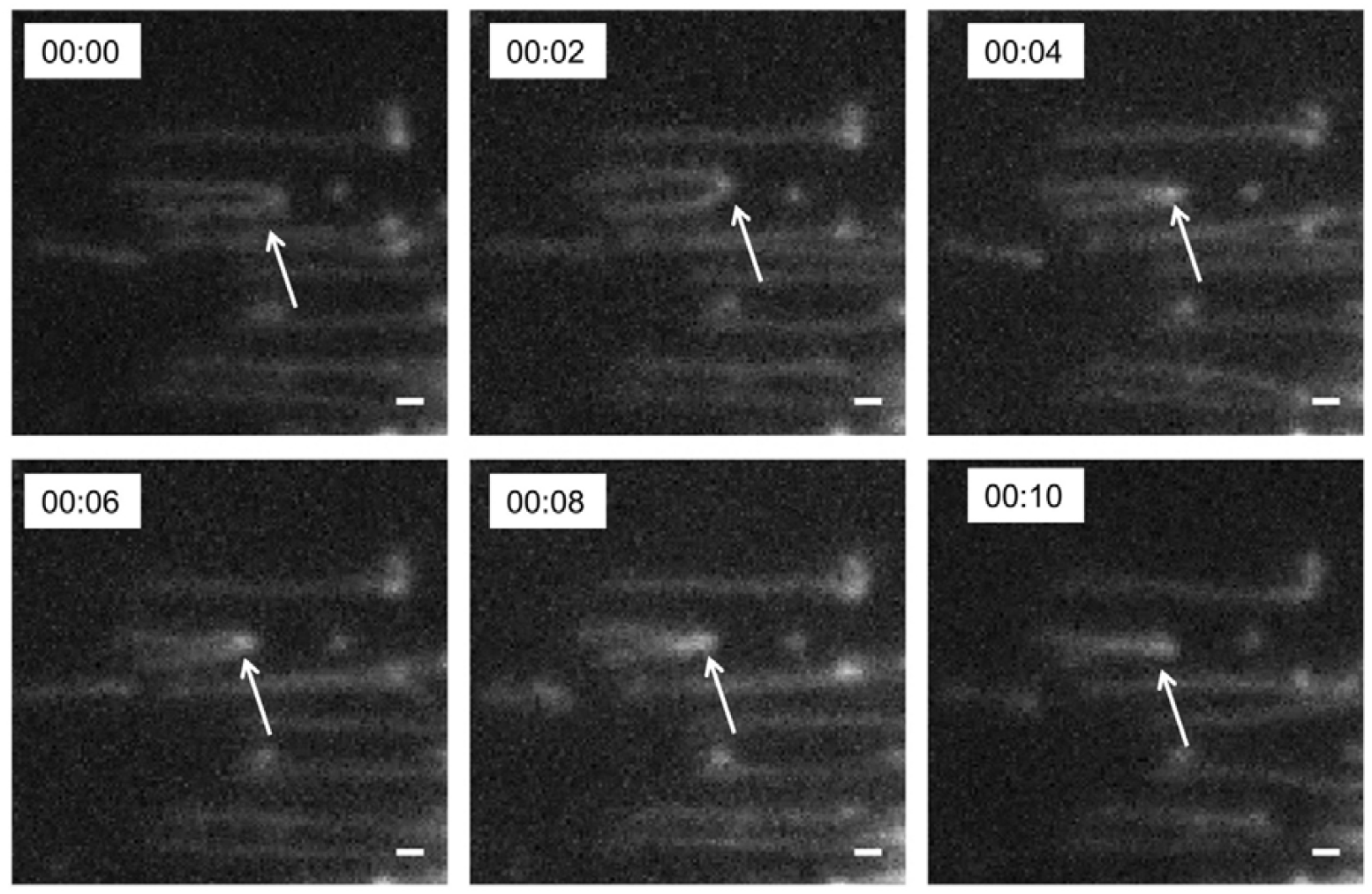
Putative DNA loop extrusion by the *U. parvum* SMC complex. Shown are sequential images of DNA molecules (linearized bacteriophage λ DNA) end-mounted on a coverslip at 2 s intervals. Arrows indicate the putative DNA loop formed by the SMC complex

### 4. *U. parvum* SMC complex is capable of loop extrusion under *in vitro* conditions

Previously, we were able to show, including in single-molecule experiments, that the Smc protein of *U. parvum* is capable of compacting DNA, and this compaction is not associated with the binding and hydrolysis of ATP [13]. However, in single-molecule experiments, in which the ability to extrude loops by eukaryotic SMC– complexes was previously demonstrated – condensins [8], cohesins [9], and Smc5/Smc6 complexes [10], *U. parvum* Smc did not exhibit loop extrusion. *B. subtilis* Smc, as well as the entire Smc-ScpAB complex, also did not exhibit such an ability. To our knowledge, the ability to extrude loops under *in vitro* conditions in single-molecule experiments has not yet been demonstrated for bacterial Smc-ScpAB complexes. In this work, we were able to demonstrate loop extrusion by the *U. parvum* Smc-ScpAB complex for the first time.

## Conclusions

1. The obtained proteins of the *B. subtilis* and *U. parvum* SMC complexes interact with each other. Smc and ScpB of *B. subtilis* form dimers, while *U. parvum* Smc forms not only a dimer, but also a putative tetramer. ScpB of *U. parvum* does not demonstrate dimerization. The interaction of *U. parvum* Smc and ScpAB is confirmed by the effect of ScpAB on the ATPase activity of Smc and its interaction with DNA.
2. The ScpAB subcomplex modulates the interaction of Smc of *B. subtilis* and *U. parvum* with DNA, weakening the formation of large complexes.
3. The *U. parvum* ScpAB subcomplex interacts with DNA.
4. The *U. parvum* ScpA protein and the ScpAB subcomplex activate the ATPase activity of the Smc protein significantly stronger than *B. subtilis* ScpAB.
5. Under the conditions of a single-molecule experiment in the presence of the *U. parvum* Smc-ScpAB complex, putative DNA loop extrusion occurs.

## Acknowledgments

The study was supported by the grant of the Russian Science Foundation No. 24-74-10022, https://rscf.ru/project/24-74-10022/. The work was carried out using scientific equipment of the Center of Shared Usage “The Analytical Center of Nano- and Biotechnologies of SPbPU”.

## Author Contributions

N.R.: Investigation, Formal analysis; A.S.: Investigation,, Formal analysis; A.K: Investigation, Formal analysis; E.Z.: Investigation, Formal analysis; M.K.: Investigation, Formal analysis; N.M.: Formal analysis; A.V.: Investigation, Formal analysis, Supervision, Writing, Funding acquisition.

